# Mechanistic Insights into Lenacapavir-Induced Off-Pathway HIV-1 Capsid Assembly

**DOI:** 10.1101/2025.08.13.670175

**Authors:** Manish Gupta, Curt Waltmann, Nadine Renner, Yihang Wang, Leo C James, David A Jacques, Till Böcking, Gregory A. Voth

**Affiliations:** Department of Chemistry, Chicago Center for Theoretical Chemistry, Institute for Biophysical Dynamics, and James Franck Institute, The University of Chicago, Chicago, IL 60637; MRC Laboratory of Molecular Biology, Cambridge, United Kingdom; EMBL Australia Node in Single Molecule Science, School of Biomedical Sciences, UNSW, Sydney, Australia

**Author notes:** **Corresponding Author:** Gregory A. Voth, Department of Chemistry, The University of Chicago, 5735 S. Ellis Ave, SCL 123, Chicago, IL 60637, **E-mail:**.

**Keywords:** HIV-1 capsid, IP6, Lenacapavir, viral assembly, defective morphologies, molecular dynamics

## Abstract

The HIV-1 capsid is a fullerene-like shell composed of hexamer and pentamer arrangements of the capsid (CA) proteins. The cone shape of the capsid is particularly important for packaging the viral genome and coordinating nuclear entry. Lenacapavir (LEN), a potent long-acting inhibitor, has been shown to disrupt capsid morphogenesis by binding at the FG-binding pocket located between neighboring CA subunits. Interestingly, inositol hexakisphosphate (IP6), a cellular polyanion, binds within the central pore of capsid pentamers and some hexamers while playing a key role in regulating the hexamer/pentamer switch. As LEN and IP6 interact with overlapping structural elements, they can compete to influence the capsid assembly pathway and outcomes. Using coarse-grained molecular simulations, we examined capsid assembly across varying IP6 and LEN conditions. Our results reveal a concentration-dependent shift in assembly outcomes: LEN accelerates hexamer assembly and reduces pentamer incorporation, leading to malformed, multilayered, or incomplete capsids. Simulations including a model for the viral ribonucleoprotein (RNP) complex further show that LEN-treated capsids frequently fail to encapsidate the RNA genome, indicating impaired maturation. Our calculations confirm that LEN impairs the formation of high-curvature CA lattice regions necessary for closure, supporting a model of off-pathway assembly as a mechanism of viral inhibition.

## Introduction

Recent advances in understanding the versatility of the HIV-1 capsid in cellular trafficking, ribonucleoprotein complex (RNP) packaging, and nuclear transport have underscored its potential as a powerful therapeutic target. While current antiretroviral therapy (ART) regimens effectively target several critical stages of HIV-1 infection ^1^, they are limited by adverse side effects, tolerability issues, and viral resistance. These challenges have prompted renewed interest in identifying alternative drug targets, among which the HIV-1 capsid has gained notable attention^2^. The HIV-1 capsid engages cellular transport machinery to traffic towards the nuclear membrane^3^. During cytoplasmic transit, the capsid serves as a nano-container to safeguard the viral genome and facilitate reverse transcription ^4^. Strikingly, recent imaging studies have revealed that the capsid transports through nuclear pore complex largely intact ^5^. Within the nucleus, stiff viral DNA strands produced during RT are believed to trigger capsid rupture, releasing the viral DNA for integration into the host genome ^6-10^. Therefore, the HIV-1 capsid presents a promising target for combating HIV-1 infection.

In the mature virion, the capsid is composed of approximately 1200 CA monomers, which are released following the cleavage of the Gag polyprotein ^11-13^. A typical mature capsid is predominantly cone-shaped and encloses the condensed viral RNA, nucleocapsid, reverse transcriptase, integrase (IN), and other proteins collectively referred to as the RNP ^14-17^. The CA monomer contains two globular domains, the N-terminal and the C-terminal domain (NTD and CTD) ^18^. Under suitable conditions, the CTD/CTD interactions induce CA dimerization and NTD/NTD interactions lead to oligomerization forming higher order hexamer and pentamer structures ^19,20^. While the hexamers have intrinsic curvature as evidenced by the formation of tubes, the pentamers create much higher mean curvature and provide the Gaussian curvature necessary for RNP encapsulation and capsid closure. Notably, exactly 12 pentamers are required for closed capsid formation. Recent biochemical and computational studies have demonstrated that cellular polyanion inositol hexakisphosphate (IP6) plays a key role in capsid morphogenesis ^15,21-23^. In the absence of IP6, CA subunits were observed to assemble into a hexameric lattice cylinder, whereas addition of IP6 restored conical morphology ^22^. Detailed structural analyses using x-ray crystallography and cryo-EM have identified IP6 densities near the R18 and K25 ring within the central pore of both hexamer and pentamer ^22,24^, further supporting this host polyanion’s role in stabilizing the capsid architecture ^25,26^. In the pentamer, IP6 favors helix 1/helix 2 packing between adjacent NTD domains triggering the folding of the TVGG motif (residue 58-61) ^23,27^. This conformational change is believed to lock the pentamer state and prevent its remodeling into a hexamer. Strikingly, the spatial proximity between the TVGG motif and phenylalanine-glycine (FG) binding pocket suggests a potential allosteric connection between IP6 and FG peptide binding. Formed during capsid oligomerization, these hydrophobic FG binding pockets recruit host factors such as CPSF6, NUP153 and SEC24C to regulate capsid cellular trafficking and nuclear entry ^28-32^. It is noteworthy that several capsid inhibitors, PF74, GS-CA1 and Lenacapavir (LEN), also target the same FG binding interface ^33^. Among these, LEN has emerged as a particularly promising candidate due to its picomolar potency against multi-drug-resistant HIV-1 strains and prolonged virological suppression in ongoing clinical trials ^34-36^. LEN has also been proposed to exhibit multimodal mechanisms of action that target multiple stages of the viral life cycle including the immature Gag lattice, capsid maturation, and nuclear entry of the capsid ^34,37-40^.

The mechanism by which LEN influences capsid formation during the late stage of HIV-1 replication is particularly critical, as this disruption occurs at clinically relevant concentrations. Consistent with this, LEN was observed to inhibit HIV-1 maturation with a half maximal effective concentration (EC_50_) of ∼ 60 pM in cell culture assays ^41^. Furthermore, LEN is proposed to impair proper virion maturation by inducing malformed and irregular outcomes during CA assembly.

However, *in vitro* experiments have shown that sub-stoichiometric levels of LEN are insufficient to fully inhibit conical capsid formation, as substantial capsid-like particles (CLPs) were still observed at low LEN levels. One possible explanation is that these seemingly intact CLPs contain defects that cause mechanical failure of the capsid during downstream activities, although this is yet to be confirmed experimentally. Additionally, little is known about the assembly kinetics and intermediate states, which can be quite complex in the presence of drug-like molecules ^42^. Importantly, IP6 and LEN have been shown to stabilize pentamer and hexamer, respectively; hence, their combined presence is expected to alter the assembly landscape. Here, we investigate the molecular origin of alternative assembly pathways and outcomes as a function different LEN and IP6 conditions.

Often, all-atom molecular dynamics (AA MD) is considered well suited for simulating the effects of small molecules on proteins and protein-protein interfaces ^43^. However, generic forcefields are unable so far to accurately model LEN interactions^44^ and custom forcefields are still under development. Moreover, even with highly accurate all-atom forcefields, atomistic MD remains intractable due to prohibitive spatiotemporal scale of the capsid assembly system. To overcome this, we use in this work coarse-grained (CG) molecular dynamics which enables access to biologically relevant scales by simulating lower resolution representations ^15,45-49^. We have simulated a wide range of conditions to understand how different LEN and IP6 conditions influence the HIV-1 capsid assembly pathway (Fig. 1a). We modeled this range of conditions to analyze their impact on assembly trajectories and capsid morphology. At low LEN concentrations, conical capsids formed but often failed to close completely, while higher LEN levels promoted defective lattices that frequently developed malformed and over assembled structures. Our simulations reveal that LEN does not directly inhibit pentamer formation but instead shifts the assembly pathway by favoring hexamer stabilization and defect incorporation. Our curvature analysis calculations further show that at high LEN ratio, the formation of properly aligned hexamer–hexamer and hexamer-pentamer interfaces become increasingly impaired, particularly in high-curvature regions leading to lattice flattening and failure of capsid closure. Additionally, we find that capsid-RNP packaging is substantially impeded in the presence of LEN, indicating that altered assembly pathways may compromise the encapsulation of the viral RNA.

**Figure 1.**
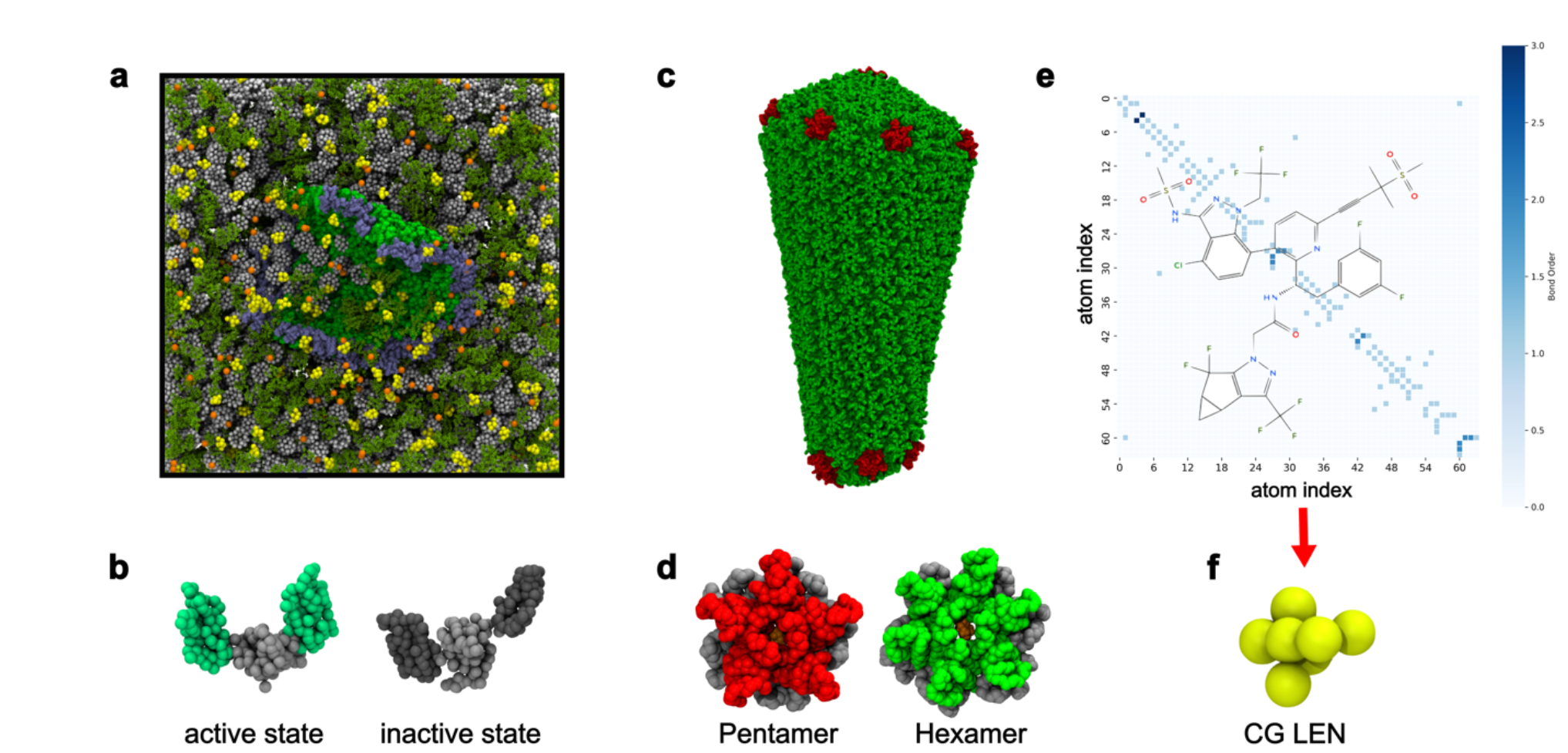
Coarse-grained (CG) model of HIV-1 CA and LEN. **(a)** Representative snapshot of the simulation setup. Free CA dimers are shown in dark green, capsid lattice is shown in green and dark gray inert crowders are shown in white, IP6 molecules in orange and LEN in yellow. **(b)** Assembly competent (left) and incompetent (right) Ultra-CG (UCG) states of CA dimer. **(c)** CG model of the full capsid made of pentamer (red) and hexamer (green). **(d)** CG model of the IP6 bound pentamer and hexamer. **(e)** Overlay of LEN molecular structure and adjacency matrix based on chemical connectivity. **(f)** CG model of LEN derived using GBCG method.

## Results

### Coarse-Grained modeling and simulation of LEN induced HIV-1 CA assembly

In solution, CA monomers and dimers exist in a dynamic equilibrium, with both forms exhibiting substantial conformational heterogeneity. Therefore, only 5-10% of the CA dimers populate an assembly-competent conformation ^20^. To simulate this behavior with our CG model, we introduced a multistate representation in which individual CA dimers called an Ultra-CG (UCG) model^50,51^ that can switch between an assembly-competent/active ([CA]_+_) or an assembly-incompetent/inactive ([CA]_-_) state (Fig. 1b). Throughout the simulation, free CA dimers underwent probabilistic switching between these two states at regular intervals, with the switching probability calibrated to reflect experimentally observed equilibrium populations. In contrast, once a CA molecule becomes incorporated into an assembling lattice, it is fixed in the [CA]_+_ state for the remainder of the simulation. Specifically, we employed a previously developed mature assembly model that was shown to reliably capture key features of capsid assembly in the presence of small molecules like IP6 and PF74 ^15,42^. In this model, 10% of CA dimers were designated as active at a switching interval of every 5×10^5^ CG time steps. This approach is a form of CG modeling in which discrete internal “states” are encoded into the low-resolution CG particles ^50^, allowing biomolecular systems to undergo state changes pertinent to conformational change or allosteric regulation.

To capture the influence of LEN, we developed a CG model guided by biochemical insights from capsid-LEN interactions. LEN is a rigid molecule featuring four ring structures (R1-R4), which form extensive hydrophobic, electrostatic and hydrogen-bonding network within the FG pocket of the neighboring CA subunits ^32,52^. R1 interacts with the NTD of both CA subunits, while R2 engages with the CA1-NTD residues. Importantly, R3 ring binds strongly with sidechains of L56, V59, M66 and other residues within the CA1-NTD hydrophobic pocket ^32^. Finally, R4 ring intercalates with CA1-NTD and CA2-CTD residues N63, M66, T169, L172, R173. The all-atom structure of LEN, derived from its co-crystal complex with the HIV-1 hexamer, was mapped onto a CG model such that each site corresponded to critical functional groups mediating CA-LEN interactions. For this, we employed the graph-based coarse-graining (GBCG) scheme, a spectral clustering algorithm designed to optimally group atoms into CG sites (Fig 1f) ^53^. The LEN molecular graph was sequentially contracted by grouping atoms with the strongest mutual connectivity, effectively preserving the key interaction motifs while systematically reducing model complexity. Attractive Gaussian potentials were used to represent interaction between LEN CG sites corresponding to R1-R4 rings and CA CG sites corresponding to residues S41, N57, Q67, K70, N74, T107, Y130, R173 to replicate binding modes observed in high-resolution crystal structures.

To examine the fidelity of our CG CA-IP6-LEN system, we begin by considering the viral capsid assembly under low IP6 and low LEN conditions (CA/IP6/LEN ratio of 8.8:1:1). Consistent with prior *in vitro* studies ^39^, CA assemblies predominantly formed tubular structures and IP6 failed to effect pentamer formation during the simulation (Fig. 2a). Pentamer defects often appear in the growing capsid lattice due to strain at high curvature regions but are seen to be transient in CG simulations the absence of IP6 or at low IP6 concentration ^15^. LEN binding promotes uniform hexamer formation and the frequency of transient pentamer formation decreased substantially. In one of the four replicas, we did not observe any transient pentamer at the growing lattice edges.

**Figure 2.**
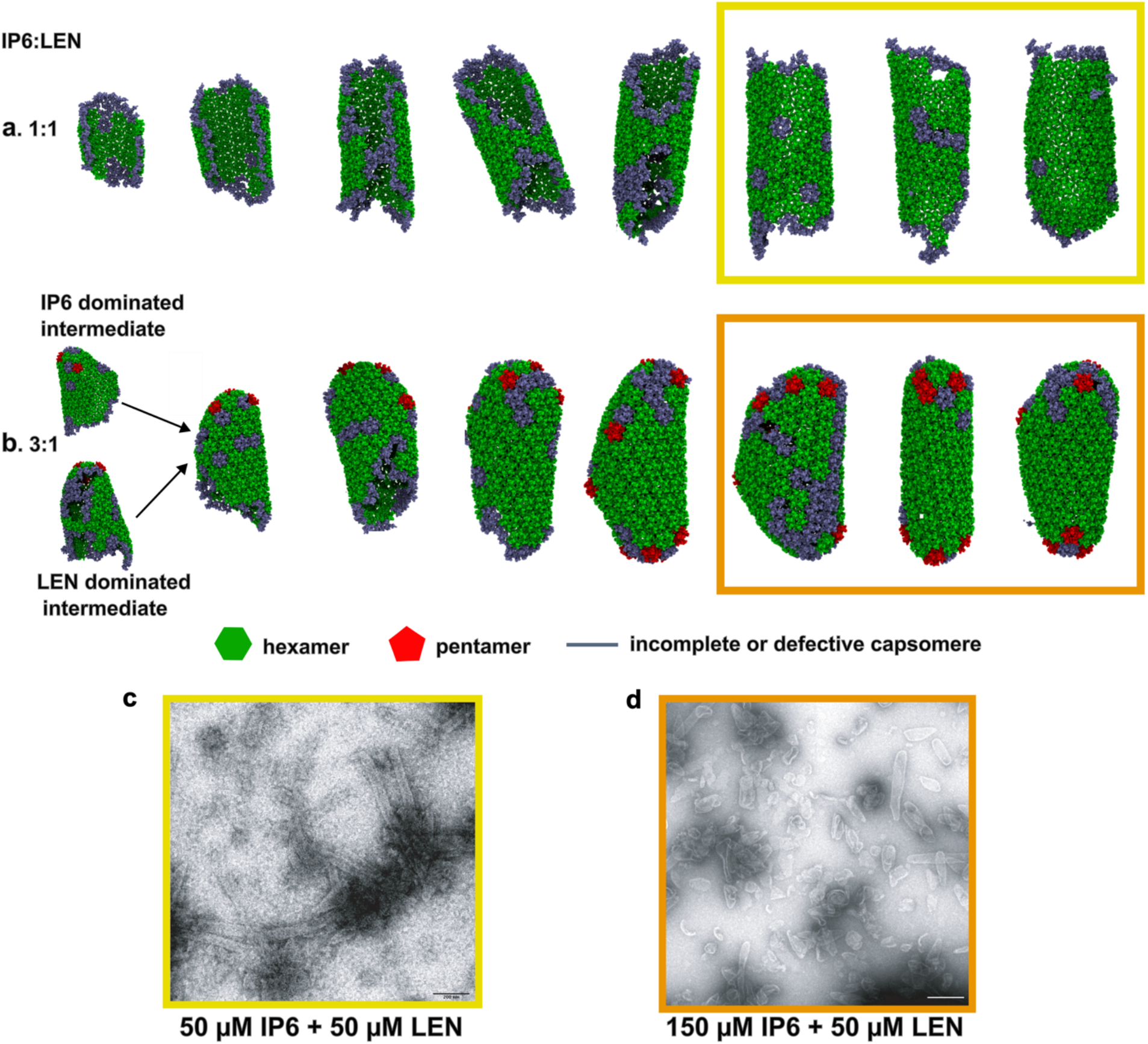
IP6/LEN competition during CA assembly. **(a)** Simulation snapshots of the capsid lattice at 50 × 10^6^, 100 × 10^6^, 150 × 10^6^, 250 × 10^6^, 300 × 10^6^ CGMD time steps under IP6/LEN ratio of 1:1. **(b)**. Snapshots of the capsid lattice at 50 × 10^6^, 75 × 10^6^, 150 × 10^6^, 250 × 10^6^, 300 × 10^6^ CGMD time steps under IP6/LEN ratio of 3:1 **(c-d) Negative stain electron microscopy images of** *In vitro* assembly products from reaction containing 75 *μM* CA, 50 *μM* IP6 and 50 *μM* LEN and 75 *μM* CA, 150 *μM* IP6 and 50 *μM* LEN. The scale bar in each image is 200 nm.

Step wise intermediates during the capsid cylinder formation are shown in Fig. 2a. Overall, capsid assembly for the IP6/LEN ratio of 1:1 was observed to be uniform, forming a curved sheet of hexamers which continued to grow in sync at both ends eventually quilting into a tubular structure. The formation of the seam region, where opposite lattice edges anneal, occurred relatively slowly. The capsid structure contained contiguous regions composed of irregular hexamers (shown in gray). Next, we experimentally reconstituted CA (75 *μM*) assembly *in vitro* in the presence of 50 *μM* IP6 and 50 *μM* LEN. We observed LEN dominated CA assembly forming tubes in this regime consistent with our computational observations and prior reports. Negative stain EM images of the capsid lattice are shown in Fig. 2c.

In contrast, IP6 has been reported to recover conical capsid formation when present at high concentration ^39,41^. Using 8.8:3:1 CA/IP6/LEN ratio, we also observed similar assembly behavior in our simulations. In this regime, capsid assembly followed two distinct pathways depending on the LEN and IP6 binding dynamics. In some cases, CA yielded a hexamer sheet akin to intermediates in the previous case owing to effective hexamer stabilization by LEN during early steps of the assembly (Fig. 2b). Next, the sheet curled into a tube-like structure. At this point, IP6 stabilized a few pentamers capping one end of the lattice while the other end continued to grow. This intermediate could potentially lead to short and capped tubes or cones observed in previous experiments in the presence of low LEN and high IP6 ^39,41^. Alternatively, IP6 sometimes was able to trap pentamer early, this forming the broad end of the capsid first (Fig. 2b). These distinct pathways likely emerge from the stochastic behavior of assembly and pentamer incorporation. In both cases, IP6 biased the assembly outcomes toward predominantly cone-shaped morphologies. Notably, high curvature regions frequently contained contiguous arrays of irregular hexamers. CA assembly in the presence of 150 *μM* IP6 and 50 *μM* LEN confirmed the emergence of both capped tubes and cones seen experimentally (Fig. 2d).

### LEN influences assembly dynamics and morphological outcomes

To better understand how different CA/LEN ratios influenced HIV-1 capsid assembly, we simulated a total of 40 replicas of capsid assembly across four different CA:LEN ratios of 8:1, 4:1, 2:1 and 1:1 at fixed IP6 concentration and compared the resulting assembly pathways and morphological outcomes. We note that 600 molecules of IP6 were added to the system (corresponding to ∼2 IP6 molecules per CA hexamer) which was found to be sufficient to promote conical assembly in previous CG simulations ^15^. Figure 3 depicts the assembly products for each of the 40 CG simulations. In total, we observed several different types of capsid morphologies: 1. canonical capsid intact and closed, 2. broken capsid with ruptured narrow or broad end, 3. defective capsid containing multiple missing CA and holes, 4. over-assembled structures, 5. Aberrant, referring to CA cluster. At CA/LEN ratio of 8:1, we observe that ∼ 50% capsids assembled cone-like outcomes while the rest assembled broken and defective capsids. In this case, CA could form pentamer state during early and late stage of the assembly process. However, the formation of all 12 pentamer was rare, as indicated by the pentamer time series profile (Fig. 4b). Instead, missing pentamer sites were occupied by irregular and incomplete capsomeres. As a result, formation of the narrow end was impaired in multiple lattices (Fig. 3a-iii, iv and x). We also observed over-assembly in one instance (Fig. 3a-ii). Similarly, at a CA/LEN ratio of 4:1, cone-shaped structures formed occasionally. Although the incorporation of a few pentamers promoted the emergence of conical geometries, regions of high curvature displayed substantial defects, including missing CA subunits (Fig. 3b-i, iv). Pentamer formation was particularly impaired, as indicated by ruptured narrow tips and persistent holes within the capsid lattice. Strikingly, in locations where pentamers failed to form, incomplete or partial hexamers were incorporated to accommodate the local curvature at both the broad and narrow ends of the capsid. Furthermore, irregular hexamers were frequently observed in high-curvature regions, especially at sites adjacent to pentamers, suggesting that the constituent CA dimers were unable to achieve the conformations necessary for proper hexamer-pentamer interface formation.

**Figure 3.**
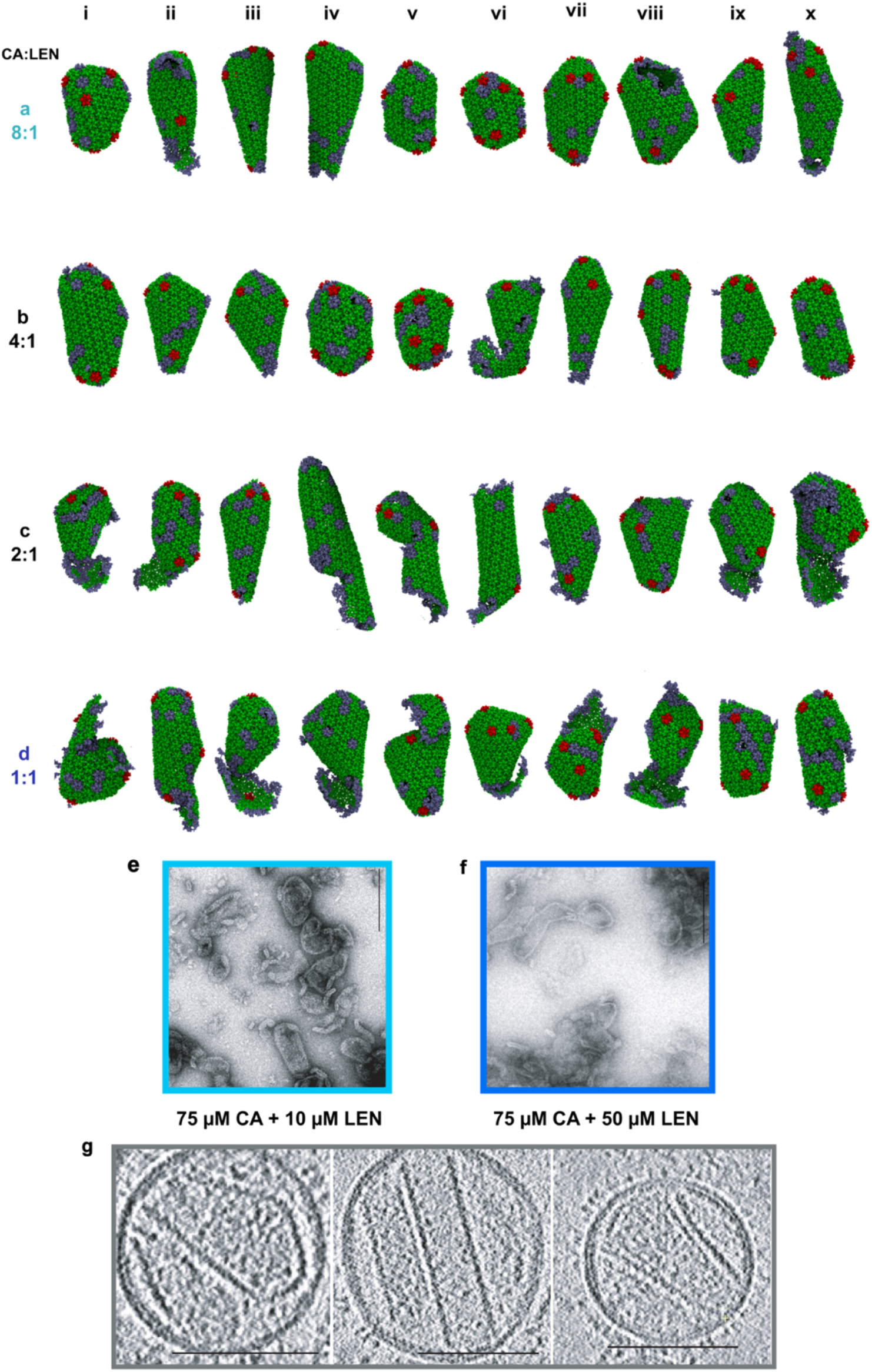
CA/LEN competition. **(a-d)** Representative snapshots of the assembly outcomes from simulations at CA/LEN ratios of 8:1, 4:1, 2:1 and1:1. Only capsid lattice is shown for clarity. **(e)** Assembly products from reaction containing 75 *μM* CA, 200 *μM* IP6 and 10 *μM* LEN and **(f)** assembly products observed in reactions with 75 *μM* CA, 200 *μM* IP6 and 50 *μM* LEN. Scale bars: 200 nm in (e) and (f). **(g)** Cryo-ET analysis of LEN-treated HIV virions (30 minutes incubation with 700 nm LEN) show over-assembled CA structures. Scale bars 100 nm.

**Figure 4.**
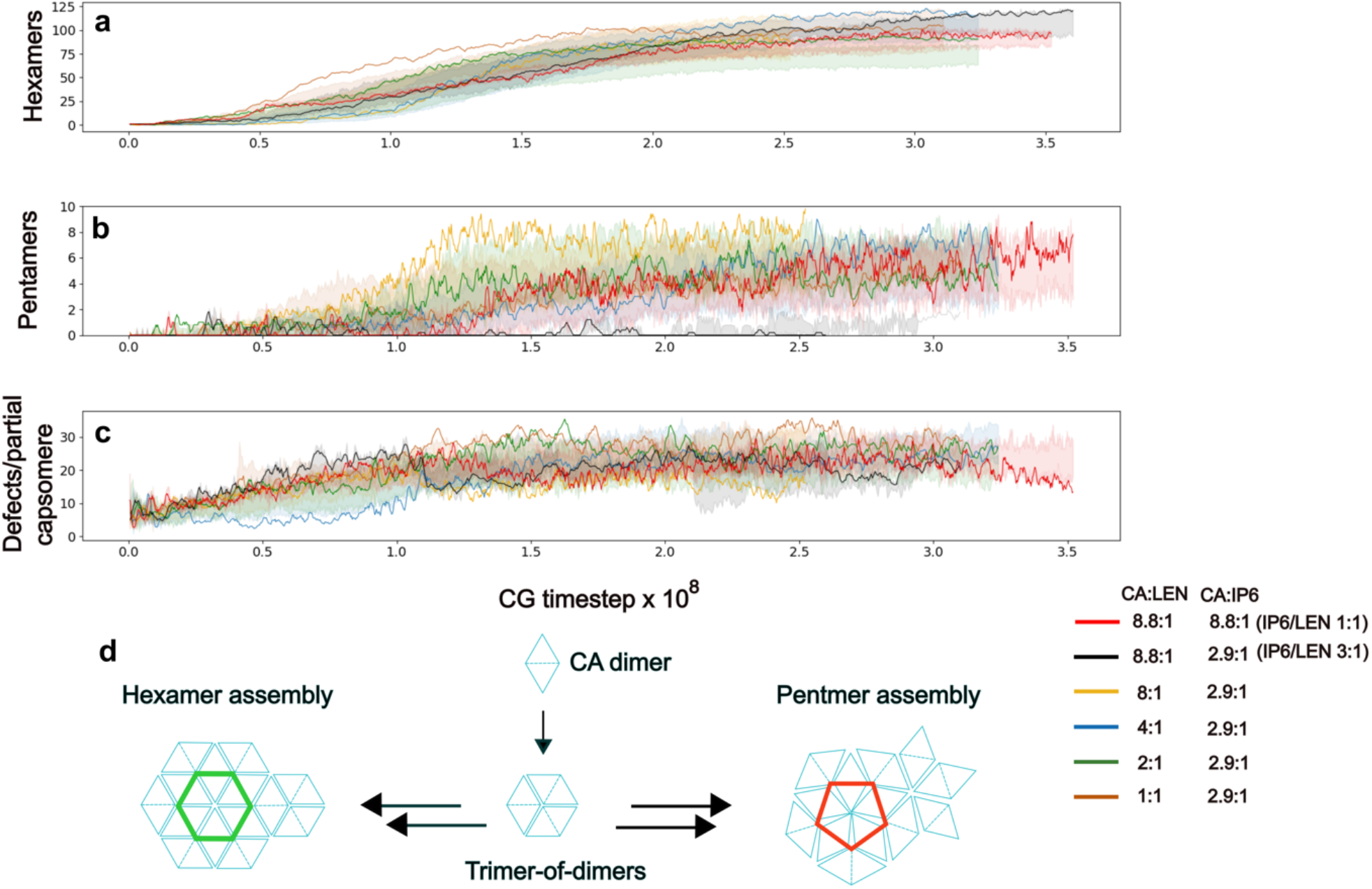
Analysis of capsid assembly dynamics. Time-series plots showing **(a)** the progression of hexamer assembly, **(b)** the total number of pentamers, and **(c)** the accumulation of lattice defects and irregular capsomeres over the course of the simulation. The shaded areas depict the standard deviation around the mean from all replica simulations and solid lines depict individual trajectories. **(d)** Schematic illustration of hexamer and pentamer assembly pathways starting from CA dimers.

**Movie S1:**
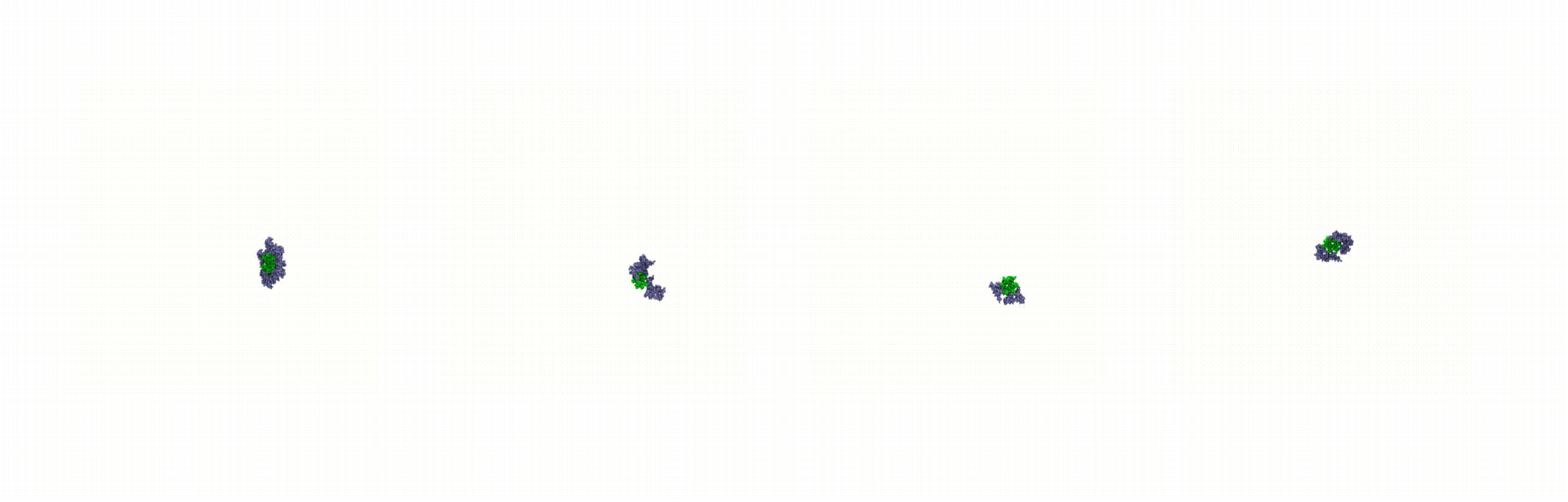
Capsid morphogenesis at low LEN levels. All four independent trajectories demonstrate that capsid can adopt conical geometry, with pentamers forming during both early and late stages of assembly. CA assembly is like WT, with the broad end forming first and growing into a hook-like structure. However, CA lattice closure is delayed and substantial defects emerge at the high curvature regions.

**Movie S2:**
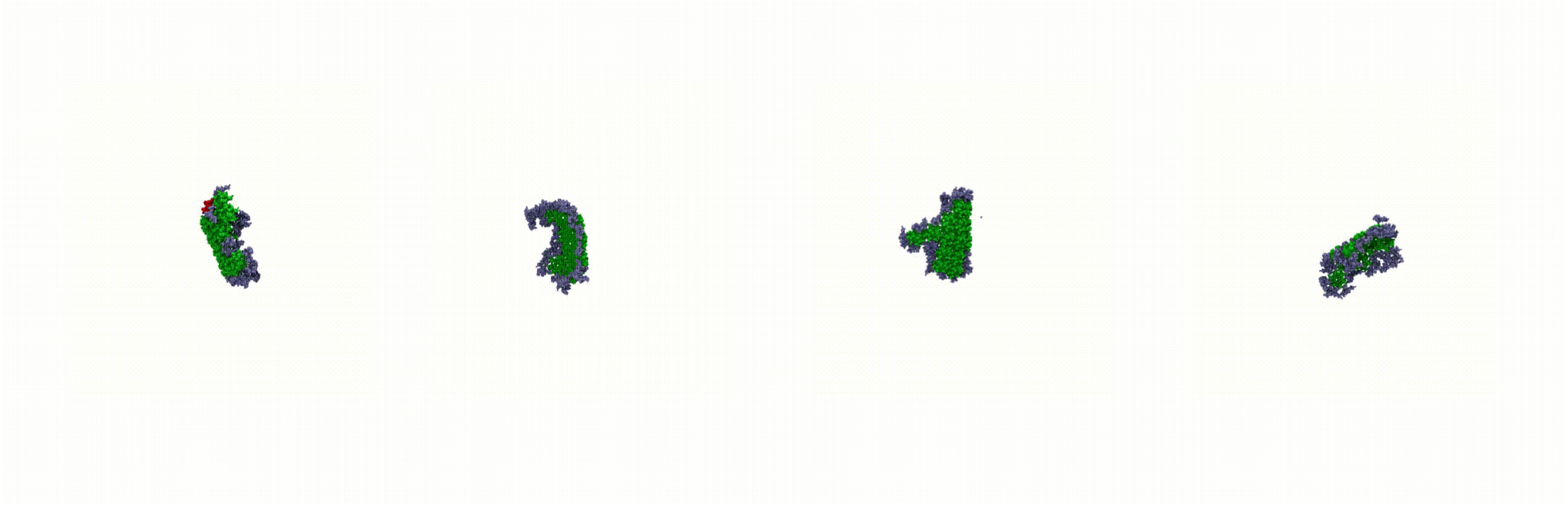
Capsid morphogenesis at high LEN levels. In one trajectory, a partially conical capsid forms, while in others, the CA lattice grows rapidly from both ends and anneals through end-to-end joining, characteristic of tubular structures. Pentamers fail to form during the late stages, preventing proper lattice closure. In all cases, an additional CA layer assembles on the lattice, reflecting elevated assembly kinetics at high LEN concentrations.

At CA/LEN ratios of 2:1 and 1:1, the distribution of capsid morphologies shifted markedly toward defective and over-assembled structures, consistent with structural and biochemical analyses of viral cores exposed to high concentrations of LEN ^39,41^. The inherent dynamic and stochastic nature of capsid assembly, modulated by the relative stabilization effects of IP6 and LEN, produced a broad array of intermediates (movies S1 and S2). In some trajectories, early trapping of pentamers by IP6 enabled the formation of a curved broad end, which under normal conditions would facilitate orderly incorporation of hexamers along the periphery, culminating in closure at the narrow tip by additional five pentamers. However, in the presence of high LEN concentration, accelerated hexamer formation disrupted this balance, leading to asymmetric lattice growth. As a result, large gaps appeared at the narrow end, which often failed to close either because one side outgrew the other or because pentamers did not form in a timely manner to generate the necessary curvature for lattice closure. These assembly defects manifested as over-assembled (Fig 3c-ii, ix, x and 3d-i-vii), cluster-like structures (Fig. 3c-v and 3d-v, viii, x) and irregular capsid shapes (Fig. 3c-i,iv and 3d-i,viii), in qualitative agreement with cryo-electron tomography (cryo-ET) images of LEN-treated virions (Fig. 3g) ^39^. It is important to note that in Fig. 3g, HIV-1 particles were first allowed to form mature virions and were subsequently treated with LEN. In contrast, LEN was present from the onset in our simulations. Nevertheless, the late-stage assembly behavior is comparable, as delayed or disrupted lattice closure allowed LEN to promote continued growth by facilitating the incorporation of remaining free CA, resulting in the formation of additional CA layers on the capsid surface. Moreover, when LEN dominated the early stages of lattice growth, a higher incidence of lattice holes and defects was observed (Fig. 3c-i, viii-x). Although native HIV-1 capsids are suggested to contain seven pentamers at the broad end and five at the narrow end, LEN-treated intermediates could still achieve substantial curvature in the absence of stable pentamer incorporation. Instead, gaps and irregular hexamers accumulated to compensate for local curvature, forming arrays of defects aligned along strain patterns previously predicted by AA MD simulations of full capsid structures ^54^. These findings suggest that LEN-induced morphologies may be susceptible to mechanical failure during later stages of the viral life cycle, including upon docking and attempted entry into the cell nucleus via the nuclear pore complex ^38^.

In summary, our simulations demonstrate that CA can assemble into capsid while adopting a varied curvature region at low LEN but it loses the ability to adopt pentamer states at high LEN, in which case hexamer formation is accelerated (Fig. 4a). Thus, the hexamer versus pentamer balance is increasingly shifted toward hexamer as LEN concentration is increased (Fig. 4a). Additionally, in some replicas, CA assembled into relatively flatter lattices (Fig. 3c-ii, iv, vi and Fig. 3d-ii, ix, x) due to delayed pentamer incorporation or a failure to accommodate the curvature required for conical geometry. A previous report proposed that LEN preferentially stabilizes the dimer interface characteristic of hexamers, but not pentamers, thereby biasing assembly toward hexamer formation ^41^. However, as shown in Fig. 4d, our simulations demonstrate that pentamers can still form from five dimers, where one monomer from each dimer contributes to the five-membered ring, collectively constituting a pentamer. This observation explains the occasional formation of conical structures, where pentamer incorporation remains feasible. More recently, structural studies have also identified the TVGG coil-to-helix switch as a key determinant of pentamer formation ^23,27,55^. It is likely that LEN binding prevents this conformational switch, locking CA into a conformation that favors hexamer over pentamer assembly.

To further validate the assembly mechanism observed in our CG simulations, we experimentally reconstituted *in vitro* CA (75 *μM*) and IP6 (200 *μM*) assembly in the presence of 10 *μM* and at 50 *μM* LEN, corresponding to CA/LEN ratios of 7.5:1 and 1.5:1 respectively. At low LEN, we observed some conical structures, whereas high LEN conditions resulted in multiple aberrant lattices (Fig. 3e, f) consistent with our simulation outcomes. Under high LEN condition, CA rapidly assembled into several open lattices, exhausting the pool of available CA before sufficient pentamers could form to close the structure. Although the CG model was originally designed to assemble only a single lattice, the over assembly behavior and delayed pentamer incorporation observed experimentally reinforce the shift toward aberrant morphologies.

### LEN induces defects at high curvature regions, favoring flatter CA lattices

To quantitatively compare the assembly pathways under different conditions, we analyzed the time series evolution of lattice curvature across all CG simulations. The CA lattice was represented as a connected graph (*G*), where each CA subunit corresponds to a node (*n*), and edges (*e*) are defined between adjacent monomers based on spatial proximity. Within the capsid lattice, edges were assigned to both intra-dimer and intra-capsomer contacts. We use the following distance criteria to construct *G*:

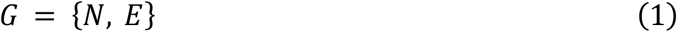

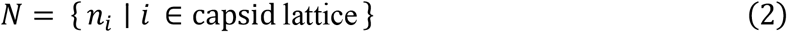

Here, *N* is the set (denoted by {}) of all nodes in the CA lattice and *E* is the set of edges spatially connecting adjacent pairs of nodes. Next, the capsid surface at each simulation frame was reconstructed by performing Delaunay triangulation over the positions of nodes in *G* ^*56*^. The local Gaussian curvature (*gc*) at each node was then computed from the angular defect of the surrounding triangles, yielding a time-resolved measure of curvature across the capsid lattice. To characterize the overall curvature distribution, Gaussian curvature values were binned and analyzed statistically. Specifically, the logarithm of tail weight (*ρ*(*gc*)) obtained from the distribution, *P(gc)*, was computed as a quantitative metric for the extent and frequency of high-curvature regions during assembly.

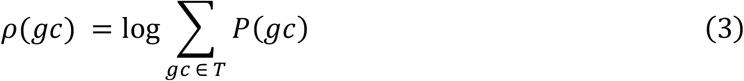

The utility of this approach is demonstrated in Fig. 5a which represents four CA lattices. In curved and conical lattices, *P(gc)* exhibits broader distribution with longer tail, corresponding to high curvature regions. Considerable density was observed for *gc* > 5 × 10^−6^. By contrast, early sheet and a tubular lattice composed of hexamers, in which CA subunits adopt relatively planar configuration, is evident by *gc* values close to zero. The resultant distribution *P(gc)* is sharply peaked between 0 < *gc* < 2 × 10^−6^. Rare minor peaks for *gc* > 10^−5^ were also observed, arising from flexible lattice edges. Therefore, *ρ*(*gc*) serves as a sensitive measure of lattice curvature, and its time evolution provides insight into the emergence of high-curvature regions during assembly. Next, we assessed the variation in lattice curvature evolution by comparing the time series profiles across all conditions (Fig. 5c). In all cases, *ρ*(*gc*) decreased initially, representing the initial nucleation of a few hexamers. At CA/IP6/LEN ratio of 8.8:1:1, the value of *ρ*(*gc*) recovered likely due to flexible CA at the lattice edges. However, the lattice curled to form a tube-like intermediate, *ρ*(*gc*) decreased sharply, consistent with a substantial reduction in the number of open edges. However, at CA/IP6/LEN ratio of 8.8:3:1 and CA/LEN ratios of 8:1 and 4:1, the *ρ*(*gc*) values shifted toward higher values (close to zero) directly preceding the incorporation of a few pentamers. During the growth phase of assembly, CA lattices grew by rapid incorporation of hexamers, leading to decrease in the fraction of high curvature CA, as reflected by decrease in *ρ*(*gc*). *ρ*(*gc*) recovered as pentamers are incorporated during later steps of assembly. At CA/LEN ratios of 2:1 and 1:1, two distinct differences were observed. First, recovery of the *ρ*(*gc*) value was slower indicating a delay in the formation of high curvature region and pentamers. Second, *ρ*(*gc*) converged at lower final values suggesting that these conditions are relatively less favorable for pentamer assembly. This indicates that CA lattices are considerably flatter in this regime. Overall, the timing and extent of curvature accumulation varied considerably, indicating that assembly kinetics and defect incorporation are sensitive to both IP6 and LEN concentration.

**Figure 5.**
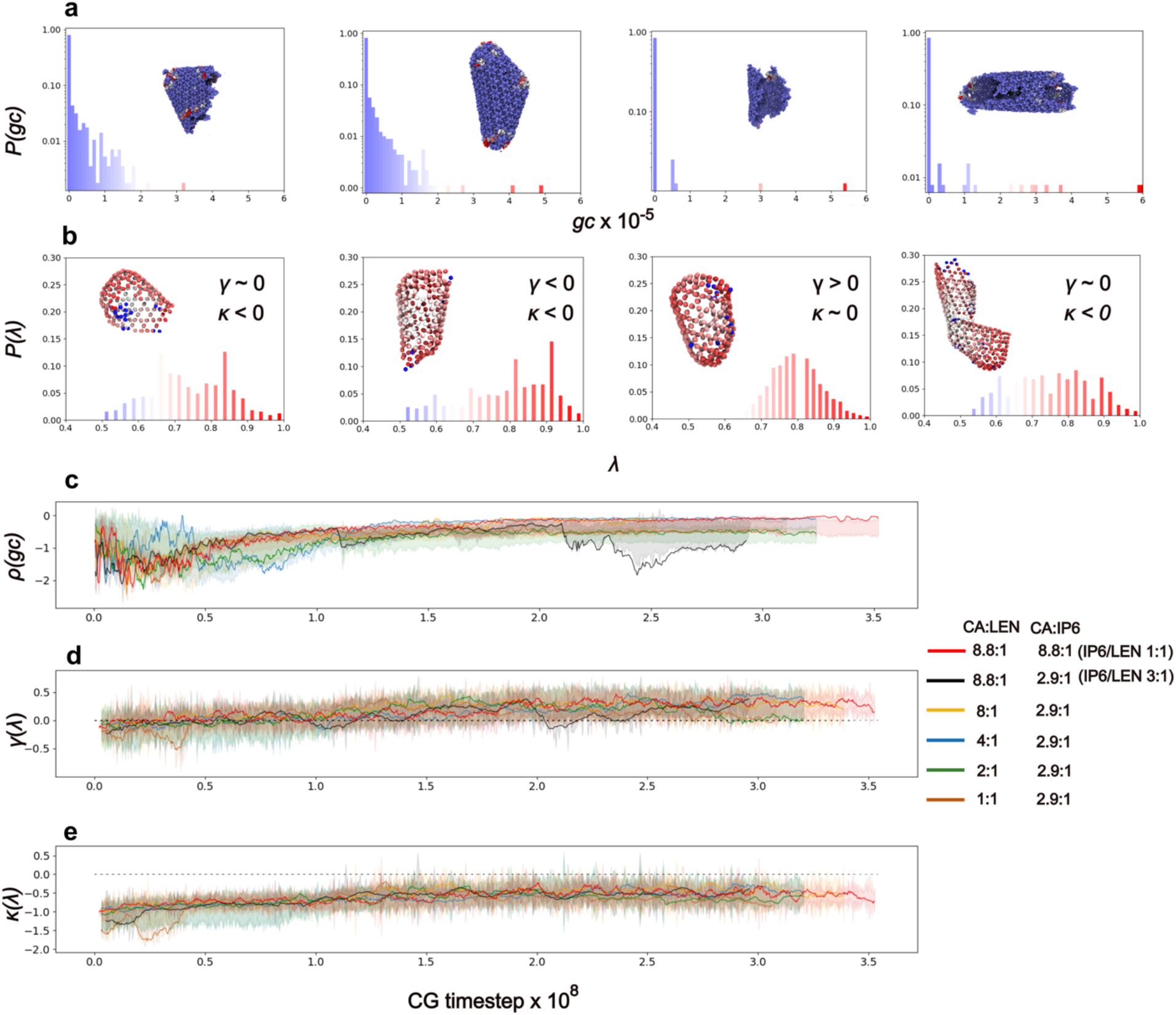
Topological analysis of the assembled capsid lattice. **(a)** We show the distribution of Gaussian curvature (*P(gc)*) for specific capsid lattices in the upper row of panels for demonstrative purpose. **(b)** Distribution of normalized eccentricity *P*(*λ*) for capsid lattice intermediates and final structures are shown for comparison. The topology of the lattice can be characterized by the skew (γ) and kurtosis (κ) of *P*(*λ*). In both (a) and (b), the color gradient of the histogram bars (blue - low values, white – intermediate and red - high values) corresponds to the range of Gaussian curvature or eccentricity and matches the color scheme used in the lattice visualizations. **(c-e)** Time series profiles of *ρ*(*gc*), *γ*(*λ*) and κ(*λ*) for different assembly conditions. The shaded regions depict the standard deviation around the mean from all replica trajectories and solid lines depict individual trajectories.

**Figure 6.**
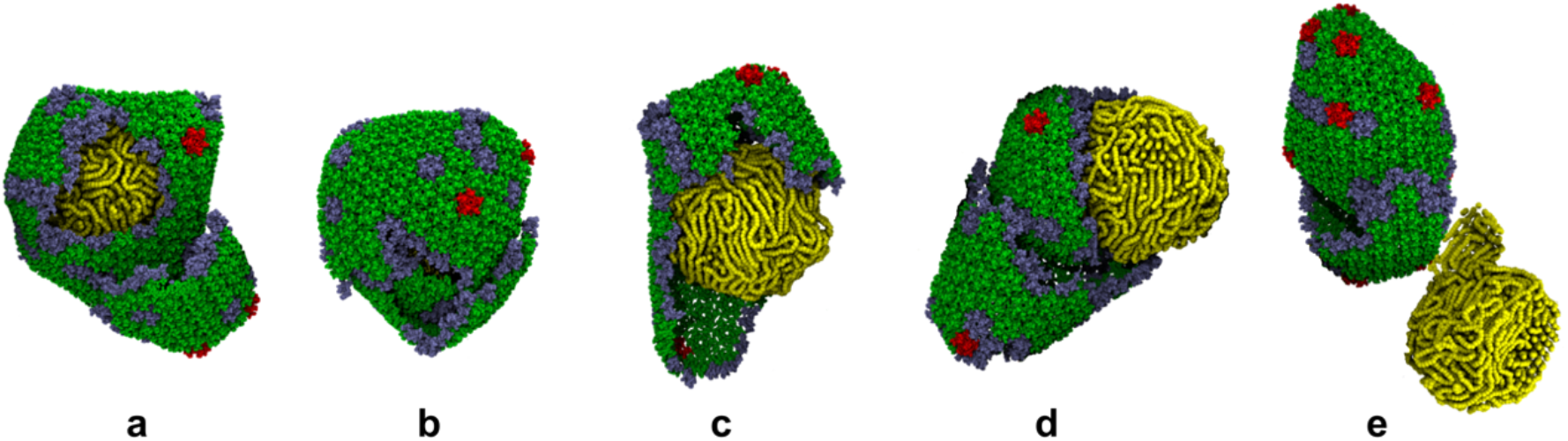
Capsid-RNP packaging in the presence of LEN. (**a-e**) Simulation snapshots showing that capsid assembled malformed architectures in the presence of LEN. LEN treated capsid failed to properly enclose RNP (yellow).

We analyzed the topological features of the capsid lattice to extract further information about the effect of LEN on capsid assembly. To evaluate how lattice morphology evolves during assembly, we computed the distribution of a local shape descriptor eccentricity, which captures the local anisotropy of the capsid lattice.

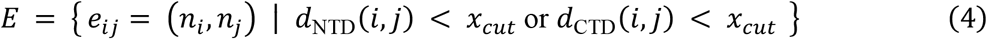

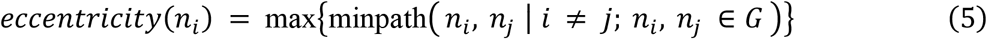

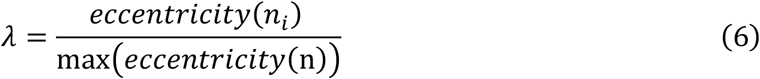

To represent the CA lattice as a spatial graph, we defined the edge set *E*. Here, *n*_*i*_ and *n*_*j*_ are nodes corresponding to CA subunits, and an edge *e*_*ij*_ is added between them if either the distance between adjacent NTD (*d*_*NTD*_) or CTD (*d*_*CTD*_) is below *x*_*cut*_. Eccentricity measures the longest of all shortest path lengths between node *n*_*i*_ and every other node in the graph G, reflecting how centrally located *n*_*j*_ is within the lattice. Finally, to normalize eccentricity values across different graph sizes or structures, we define the normalized eccentricity *λ* of a node as the ratio of its eccentricity to the maximum eccentricity observed in the graph. The eccentricity distribution *P*(*λ*) serves as an effective topological metric for capturing structural progression and anisotropy in the capsid lattice. In Fig 5b, we show representative snapshots of capsid intermediates to demonstrate the utility of this metric.

In the first panel, the distribution is roughly isotropic and broad (*γ*(*λ*)∼0, κ(*λ*) < 0), reflecting the presence of an open and minimally anisotropic lattice with a few protruding edges. These protrusions are evident in the molecular rendering, where several vertices exhibit high normalized *λ* (colored red), indicating their distal placement within the growing structure. These dendritic edges are likely to form pentamers or defects to anneal the lattice. The second panel shows a larger and more elongated lattice (κ(*λ*) < 0), with a strong left-skew (*γ*(*λ*) < 0) in *P*(*λ*). This signifies enhanced anisotropic growth, particularly along one axis, where the highest *λ* values are again concentrated near dendritic termini. These protrusions emerge during late stages of assembly and are indicative of the onset of capsid curvature or incomplete closure event. In contrast, the third panel presents a near-closed, oblong lattice with a bell-shaped *P*(*λ*), as many nodes now reside within an enclosed topology. The reduced prevalence of dendritic high *λ* values and the emergence of a peak at intermediate *λ*∼0.7 suggest a shift toward a topologically interior-dominant structure (*γ*(*λ*) > 0, κ(*λ*)∼0), characteristic of a nearly complete capsid. Finally, in the fourth panel, a broad (κ(*λ*) < 0) and roughly symmetric *P*(*λ*) distribution is observed, and all edge vertices are located at comparable distance from the center (*γ*(*λ*)∼0), suggesting the presence of an open cluster-like or over-assembled structure during late stages. The edges in this structure are uniformly spaced, eliminating localized eccentric features and results in a more balanced distribution of *λ* values across the lattice. These profiles reinforce that shifts in skewness and the spread of *P*(*λ*) are tightly linked to the geometric progression of capsid lattice. Overall, by tracking the skewness and kurtosis of *P*(*λ*), we can infer the degree of symmetry in lattice growth.

At CA/IP6/LEN ratio of 8.8:1:1, the CA lattice appears largely flat and symmetric around its sixfold axis (Fig. 5.d-e), forming a mostly uniform two-dimensional sheet during early steps of the assembly process. Correspondingly, *P*(*λ*) follows a near-normal distribution; *γ*(*λ*)∼0 values represent uniform growth of the hexamer lattice. As the lattice folds into a tube (Fig. 5d), some jagged edges disappear healing the lattice such that *γ*(*λ*) increases toward values mildly greater than 0. However, *γ*(*λ*) value again decreases to values close to 0 due to irregular growth at open ended edges. In contrast, *γ*(*λ*) values were observed to be consistently greater than 0 indicating the restoration of closed capsid at CA/IP6/LEN ratio of 8.8:3:1. Similarly, under low LEN concentration (CA/LEN ratios of 8:1 and 4:1), the capsid lattice grew relatively uniformly and incorporated pentamers frequently as evidenced by early onset of *γ*(*λ*) > 0. However, κ(*λ*)∼0 suggests that the capsid lattice did not always close properly during late steps of assembly. At high LEN (CA/LEN ratios of 2:1 and 1:1), the capsid lattice exhibited anisotropic growth as indicated by *γ*(*λ*) < 0 and κ(*λ*) < 0 during the early steps of the assembly process. The capsid lattices heal by incorporation of pentamers or defective capsomeres resulting in gradual rise in *γ*(*λ*). However, *γ*(*λ*) values decrease again due to onset of the anisotropic edges during growth phase of assembly. The capsid lattice failed to incorporate sufficient pentamers to anneal the lattice instead continues to grow resulting in over-assembly as indicated by *γ*(*λ*) close to 0. Across varying LEN concentrations, the assembly trajectories of the capsid lattice diverged subtly yet meaningfully. In all cases, the skewness *γ*(*λ*) remained close to zero during early stages, indicating that LEN promotes largely isotropic assembly biased toward hexamer formation. However, the influence of IP6 became evident at low LEN levels, where anisotropic lattice edges heal by pentamer incorporation, facilitating partial or full lattice closure. In contrast, high LEN concentrations promoted anisotropic and overextended lattice morphologies, indicated by initial negative skewness and kurtosis. These conditions impaired timely pentamer incorporation and led to persistent open edges, ultimately favoring over-assembly. Thus, different LEN concentrations were found to steer the capsid lattice along distinct assembly trajectories by modulating the balance between curvature formation and anisotropic growth.

### LEN impairs efficient viral RNP packaging

In mature virions, proper encapsulation of the genomic RNA complex is essential for a productive infection. Although the molecular details of RNP packaging by CA remain poorly understood, this process is proposed to proceed via nucleation of capsid lattice around the condensed RNP complex. The capsid lattice gradually grows into a conical capsid with RNP often localized at the broad end. Notably, the mature capsid must contain sufficient empty space to support dsDNA synthesis during reverse transcription ^6^. Therefore, LEN driven capsid assembly may disrupt virion maturation through multiple mechanisms. First, accelerated lattice growth may compromise proper RNA encapsidation, potentially exposing the viral genome to cytoplasmic immune sensors or interfering with reverse transcription, which requires an intact capsid as a reaction compartment. Second, the formation of multi-layered capsids could physically limit the available space for reverse transcription. Third, aberrant assembly may render the capsid unstable, leading to premature uncoating or impaired nuclear import. To examine how LEN affects RNP encapsidation, we simulated a capsid–RNP composite system (see Methods). In the absence of LEN, capsid assembly followed the canonical pathway, enclosing the RNP at the broad end of the cone. In the presence of 4:1 CA/LEN, capsid could enclose the RNP but the resulting structures deviated substantially from canonical morphologies. The RNP remained broadly localized near the wide end, but the surrounding lattice frequently formed broken, defective or multi-layered structures, including multiple incomplete shells around the RNP. In a subset of simulations, the capsid failed to enclose the RNP. Given the structural and functional advantages conferred by the conical geometry during intracellular trafficking and nuclear import, these LEN-induced morphologies likely correspond to noninfectious or severely attenuated viral forms.

## Discussion

HIV-1 therapeutic strategies have advanced substantially through efforts to elucidate and target key biochemical processes underlying viral replication. Among these, the capsid inhibitor LEN has emerged as a long-acting agent with picomolar potency. Notably, LEN exhibits antiviral activities at both early and late stages of HIV-1 replication. During early steps of HIV-1 replication, LEN is believed to hyper-stabilize the preformed capsid ^38^, leading to premature uncoating or impaired nuclear import. In addition, LEN is also believed to interfere with capsid maturation during late stages of viral replication ^41^. During clinical trials, LEN is administered twice annually, maintaining plasma concentrations in the low nanomolar range for several months. Thus, the picomolar activity of LEN is important in achieving long-acting antiviral efficacy.

In this study, we investigated the molecular mechanisms by which sub-stoichiometric concentrations of LEN alter the HIV-1 capsid assembly pathway, leading to non-productive assembly outcomes in a majority of cases. LEN has been hypothesized to favor the hexamer state and block pentamer assembly. However, it has not been clear how LEN inhibits pentamer assembly without specifically binding with pentamer ^24^. In fact, curved and conical capsid lattices, which likely require pentamers, have been observed even in the presence of LEN. Contrary to prior suggestions that LEN-stabilized dimers impede pentamer formation due to an incompatibility with the odd-numbered architecture ^41^, our findings support a model in which CA dimers can indeed participate in pentamer assembly. Specifically, each dimer contains two CA domains that can be shared between adjacent pentamers and hexamers, allowing pentamers to assemble directly from CA dimers. CA dimerization typically occurs through the CTD/CTD dimer interface ^57,58^, i.e. the NTD/CTD pocket required for LEN binding is not formed during dimerization. Instead, this pocket emerges during higher-order oligomerization. Taken together, these observations support a model in which CA first oligomerizes, followed by LEN binding that locks the TVGG switch in the hexamer state. This mechanism provides a more consistent explanation for how LEN promotes hexamer assembly without directly blocking pentamer formation.

We have also demonstrated that LEN binding enables alternative pathways that favor non-canonical intermediates and aberrant morphogenesis. LEN did not suppress pentamer formation; rather it accelerates hexamer assembly, perturbing the hexamer-pentamer balance necessary for capsid closure ^15^. High curvature regions of the capsid are then geometrically constrained forcing hexamer-hexamer and hexamer-pentamer pairs to adopt high tilt conformations. In the presence of LEN, hexamers often failed to adopt these high tilt states resulting in irregular hexamers and holes at the high curvature regions. Hexamers adjacent to pentamers were also often malformed. We note that structural characterization of hexamer-hexamer interfaces has identified a ‘tilt’ switch motif that regulates the relative orientation of adjacent hexamers ^55^. We therefore propose that LEN binding can influence this tilt switch, favoring flatter configurations and thereby contributing to lattice defects in high curvature regions. Further AA MD simulations of LEN-bound capsid will be necessary to more completely evaluate the effect of LEN on both hexamer-hexamer and hexamer-pentamer interfaces. Relatedly, FG repeat peptides such as CPSF6, SEC24C and Nup153, which also bind to the same FG binding hydrophobic pocket, have already been shown to influence hexamer tilt states. CPSF6 and SEC24C bound hexamers favor low tilt conformations and Nup153 bound hexamers prefers the high tilt state ^55^. Therefore, structural studies focused on the trimer interface of LEN-treated capsid could provide valuable information about the molecular mechanism underlying capsid defect formation.

In mature virions, occasional formation of intact capsids under sub-stoichiometric LEN might appear inconsistent with the ultrapotent activity of the inhibitor. Our findings suggest that capsids assembled under these conditions appear intact but exhibit substantial structural defects, particularly at the narrow end, which could predispose them to mechanical failure at later stages. Allosterically correlated strain patterns are inherently present on the capsid surface in the presence of the RNP and other host proteins ^54^. In the LEN-induced CG assemblies, we observed similar striations on the capsid surface, accompanied by lattice defects, small cracks, and irregular hexamers. These features resemble crack propagation events reported during endogenous reverse transcription. To relieve strain, the lattice incorporated structural irregularities. We speculate that such morphologies may compromise the capsid’s ability to protect the viral genome and increase susceptibility to premature uncoating. While direct simulation of capsid rupture in the presence of LEN is computationally prohibitive at present and beyond the scope of this study, our findings point toward a possible failure mechanism in LEN-driven capsid assemblies.

There is growing evidence that HIV-1 capsid flexibility is essential for engaging host factors and facilitating nuclear import ^46^. LEN, by contrast, has been reported to hyper-stabilize the capsid, reducing its intrinsic flexibility. We note that our simulations have not considered the explicit effect of host factors which interact with the HIV-1 capsid using the same FG binding pocket for downstream activities. LEN-treated capsid was shown to dock at the nuclear envelope, presumably via interaction with the cyclophilin-homology domain of Nup358, but failed to enter NPC channel ^38,40^, presumably because the FG binding sites occupied by LEN cannot interact with the FG motifs of nucleoporins. Conversely, the FG motifs of host factors in the cytoplasm, at the nuclear pore complex and inside the nucleus may interfere with LEN binding at its target pocket highlighting a far more complex and dynamic environment, which remains the subject of ongoing investigations.

## Materials & Methods

### Coarse-grained (CG) model details

In this study, LEN was coarse-grained using the graph-based CG method GBCG. In this approach, the atomic-level connectivity matrix of LEN, derived from its bonded interaction network, was subjected to spectral decomposition. The resulting eigenmodes were clustered to identify chemically and dynamically coherent domains, which were used to define a 7-site CG representation of the molecule. To model CA-LEN binding, a minimal interaction potential was implemented between specific sites on the CG LEN particle and the pre-assembled CG hexamer interface. The interaction sites were selected based on known structural binding locations of LEN within the FG pocket of the CA hexamer. These interactions are given by

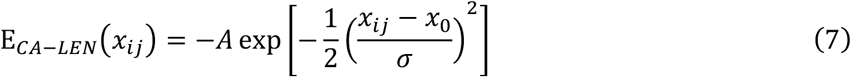

Here, *x*_*ij*_ is the separation distance of CG sites and *x*_0_ is the location of energy minima determined from the co-crystal structure of LEN bound to HIV-1 capsid hexamer (PDB 6VKV) ^52^. The parameters *A* and *σ* describe the depth and width, respectively, of a Gaussian energy well used to model LEN’s interaction within the capsid binding pocket.

We modeled the viral RNP complex as 2 polymers, which are each 3000 beads in length (the first 112 are of a different type to represent the 5’ UTR), and 35 integrase tetramers. Each polymer bead represents 3 bases and bound nucleocapsid protein in the dimeric RNA genome which is approximately 9000 base pairs long. Each bead has a diameter which is enforced by a soft exclusion potential shown below in Eq. 10, where A is 10 kcal/mol and *x*_*cut*_ is 3 nm. Consecutive CG particles, regardless of type, were connected by harmonic springs to maintain polymer continuity, with the bonding potential defined as:

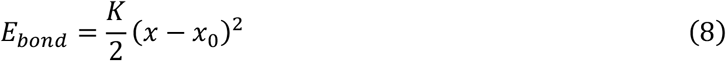

where *x* is the separation distance, and *x*_0_ = 1.0 *nm* is the equilibrium bond distance. The bond coefficient is K = 10 kcal/mol/*nm*^1^. We also apply these bonds to the secondary structure contacts found in the 5’ UTR according to the map from Goodsell et al ^59,60^. This includes the contacts between the two different chains in the dimerization sequence of the RNA. We also add angle potentials.

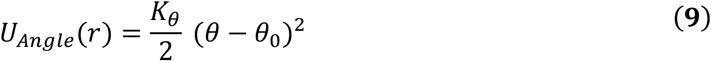

To prevent unphysical bending of the RNA chains. The equilibrium angle (*θ*_*_) is completely straight, 180°, although the bending constant is relatively weak, 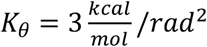. We use a gaussian attraction to model the condensation of the RNA globule due to electrostatic interactions between NC and the RNA as in Eq.12. Differing attraction strengths are used between the structured 5’ UTR and the rest of the RNA polymer (see Supplementary Materials for details).

The integrase tetramers consist of 4 beads each representing one integrase unit. They are bonded together and to the RNA/NC using harmonic bonds of the form shown in Eq. 8, where 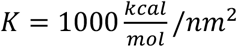 and the bond lengths vary, see Supplementary Materials. Interactions between the integrase units and RNA/NC beads are given by a gaussian attraction shown in Eq. 12. Here the width, *σ* = 0.5 *nm*, the equilibrium distance,*x*_0_ = 4 *nm*, and the interaction strength, 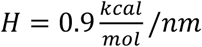, which was calibrated to control the amount of surface accessible integrase that CA can interact with (see Supplementary Information).

Experiments have suggested that integrase is necessary for proper encapsulation of the RNP by the growing capsid and thus we place attractive interactions between CA and the integrase while only volume-excluding interactions exist between CA and the rest of the RNP ^61^. Interactions between CA and IN are applied through each of the 4 integrase sites on the tetramer and site 115 on each CA. Interactions between CA and the RNA/NC are given by soft repulsions as in Eq. 10, where the diameter, *x*_*cut*_ = 2 *nm*. Attractive interactions come from a Gaussian potential in Eq. 12 where *σ* = 0.2 *nm x* = 3 *nm*, and 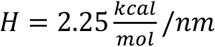. Interactions between the other RNP beads and CA are left as a soft repulsion with a cutoff, *x*_*cut*_ = 3 *nm*. These interactions and interactions between RNA/NC and the integrase were calibrated together to ensure successful assembly and RNP encapsulation in the absence of LEN, see Supplementary Information for details.

We used a previously developed solvent-free CG model of HIV-1 CA and IP6 ^15,48^. Here, we briefly describe the key features relevant to our simulations. Each monomer was represented by two independent elastic network models (ENM) for the NTD and CTD domains of a coarse-grained CA monomer, collectively comprising 130 Cα atoms corresponding to α−helices of the CA subunit. The CG representation of the NTD was constructed using Cα positions from PDB structure 3H4E ^62^. The potential energy of the bonds (*E*_*bond*_) was given by equation 8. To preserve the local secondary structure of the CA subunit, CG particles within each α−helix were connected using harmonic springs. Additionally, to preserve the relative orientation of the α−helices, each CG particle was connected to its nearest neighbors from other helices as well. A weak ENM was used to preserve inter-domain connectivity. CA CTD was modeled from PDB 2KOD using a similar approach ^63^. CA dimers were then formed by connecting the CG particles representing CTD/CTD interface of two CA subunits. Note that our primary objective was to maintain the overall molecular shape of CA dimers, rather than to reproduce atomic level details as this is a CG model.

To represent macromolecular crowding, an inert Ficoll70 molecular crowding was used ^64^. Ficoll70 was represented by two CG spheres, each of which consisted of 42 CG beads and ∼5 *nm* diameter with excluded volume separation of 1 *nm*. Excluded volume interactions were modeled using a soft cosine potential.

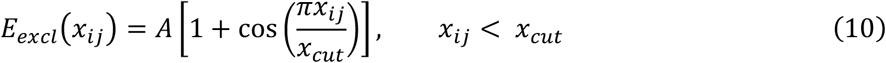

Here, *x*_*ij*_ is the separation between CG particles and *x*_*cut*_ = 10Å the onset of excluded volume repulsion for CG sites *i* and *j*. The value of *A* is 10 *Kcal mol*^−1^Å^−1^. This ensured CG particles have an effective radius and prevented any unphysical overlap between them.

To capture key intra-capsomere interactions relevant to CA assembly, inert “ghost” particles were embedded into the CG structure of CA at selected positions; residues 51, 57 and 63 on the NTD and residue 204 on the CTD. These ghost sites facilitated assembly by interacting exclusively with designated binding interfaces on adjacent CA subunits (assembly competent) when active and remaining inert when inactive (assembly incompetent).

The interaction between ghost and specific non-ghost CG particles was governed by a tunable attractive potential defined by a double-Gaussian function:

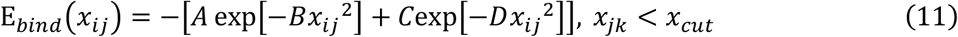

Here, *x*_*ij*_ is the separation between CG particles *i* and *j* with a cutoff radius *x*_*cut*_ = 25Å. The parameters *A* and *B* describe the depth and width of short-range CA/CA interaction. Similarly, the parameters *C* and *D* represent the depth and width of long-ranged component of Gaussian well. In our simulations, *A* = 1 *Kcal mol*^−1^, *B* = 0.1 Å^−2^, *C* = 2 *Kcal mol*^−1^ and *D* = 0.01 Å^−2^. Further details on the CG models for CA and the crowding agents are provided in previous CG studies ^15,48^.

IP6 was modeled using a single-site CG particle placed at the center of mass and the interaction between IP6 and the CA was modeled using a single-well Gaussian potential:

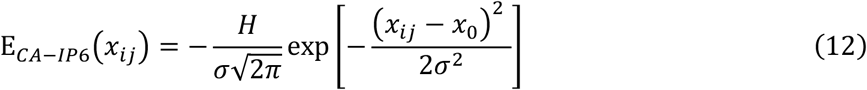

where *x*_0_ = 8.3 Å corresponds to the average distance from the center of the R18 ring to the IP6 binding site, as determined from preassembled capsid structure (Fig. 1). All CG models and simulation inputs were prepared using Moltemplate ^65^.

### CGMD simulations

Self-assembly of HIV-1 CA was simulated using CG MD in the canonical (constant NVT) ensemble at 300 K with Langevin thermostatting ^66^ and a damping time of 100 ps under periodic boundary conditions. For efficient sampling of nucleation events, a swarm of 100 parallel simulations, each initialized with randomly distributed components, was run for 10^7^τ. Trajectories yielding a nucleated cluster (fewer than 8 CA dimers) were selected to seed production simulations. System coordinates were recorded every 5 × 10^5^τ for post-processing. In the primary setup, a total of 880 CA dimers and 2500 inert crowding agents were randomly placed inside a 90 × 90 × 90 *nm*^3^ simulation box. For 1:1 IP6/LEN condition, we randomly added 200 molecules of IP6 and LEN each and performed four independent replicas of this system for 3 × 10^8^ CGMD steps. Similarly, IP6/LEN ratio of 3:1 was constructed by placing 600 molecules of IP6 and 200 molecules of LEN randomly in the simulation box. To investigate how varying CA:LEN ratios affect assembly dynamics, systems containing 880 CA dimers were simulated with 220, 440, 880 or 1760 LEN molecules, corresponding to CA:LEN ratios of 8:1, 4:1, 2:1, and 1:1, respectively. Each condition included 600 IP6 molecules and was run with 10 independent replicas for 2.5 × 10^8^ − 3.5 × 10^8^ CGMD time steps or until more than 80% free dimeric CA pool was exhausted.

For capsid-RNP simulations, a separate system containing 800 CA dimers, 400 LEN, 600 IP6 and 2000 inert crowders was constructed in a 88 × 88 × 88 *nm*^3^ volume; 10 replica trajectories were simulated. Prior parameterization of this model identified a switching interval of 5 × 10^5^ τ as suitable for generating mature capsid-like structures at a representative fraction (∼10%) of assembly competent species. Note that the estimated local protein concentration in HIV-1 virion is approximately 4 mM. After proteolytic cleavage, around 3000 CA monomers are released within a virion volume equivalent to a sphere of 140 nm diameter. Our system contains 800 dimers (1600 monomers) in a 88 *nm*^3^ box, which is about half the volume of a virion and CA contentAll CGMD simulations were carried out using LAMMPS version 21Jul2020 ^67^.

### Data analysis and visualization

Graph-based analysis of assembled CA lattices was conducted using the NetworkX 2.1 Python package (http://networkx.github.io/), based on two proximity criteria. Inter-hexameric connections were identified through CTD dimer interfaces, defined by CG type 102 pairs within 2 nm. Intra-hexameric contacts were assigned when CG type 8 particles, corresponding to the NTD helix 1 position, were within 1.5 nm of each other. Time series of cluster properties were analyzed and plotted using Matplotlib ^68^. Visual representations of selected structures were rendered with VMD 1.9.3 ^69^. To quantify lattice curvature during assembly, Gaussian curvature was computed from the reconstructed capsid surface using Delaunay triangulation over center of mass positions of each CA monomers. The angular defect at each vertex provided local curvature values, which were binned to generate Gaussian curvature histograms for each simulation frame. The relative weight of the histogram tail was used to track the time series profiles of high-curvature regions over time for each condition. In addition, topological eccentricity was calculated from the graph network of each assembled lattice. Eccentricity values were binned to obtain framewise distributions, and statistical moments (skewness and kurtosis) were computed to characterize the evolving structural asymmetry. Time series of these metrics were used to characterize lattice organization and to distinguish between isotropic and anisotropic growth behaviors.

### Negative staining electron microscopy (EM) of CA structures self-assembled in the presence of IP6 and LEN

Recombinant CA was expressed in *E. coli* C41 cells. Cell lysate was clarified by centrifugation. CA in the supernatant was precipitated with 25% ammonium sulphate, collected by centrifugation, resuspended, and dialysed against 50 mM MES (pH 6.0) containing 20 mM NaCl and 1 mM DTT. CA was purified using cation exchange chromatography with a gradient from 20 mM to 1 M NaCl followed by size exclusion chromatography with Tris (pH 8.0) containing 20 mM NaCl and 1 mM DTT and snap frozen for storage until use in assembly reactions.

Thawed CA was dialysed against 50 mM MES (pH 6.0) containing 1 mM DTT. CA assembly was carried out at a final concentration of 75 µM in the presence of 2% DMSO. LEN and IP6 were added to the specified concentrations to initiate assembly, and the reaction mixture was incubated overnight. A sample from the assembly reaction mixture (5 µL) was applied to a carbon coated grid (Cu, 400 mesh, Electron Microscopy Services) that had been cleaned by glow discharge. The grids were then washed and stained with 2% uranyl acetate. Micrographs were acquired using a Tecnai Spirit (FEI) operated at 120 keV and recorded with a Gatan CCD camera. Images were collected with a total dose of ∼30 e^-^/Å2and a defocus of 1–3 µm.

### Cryo-electron tomography of LEN-treated HIV particles

Replication deficient VSV-G pseudotyped HIV-1 virions were produced in HEK293T cells using pCRV1-GagPol, pCSGW and pMD2.G, harvested at 24–48 h post transfection, filtered (0.22 μm) and concentrated by ultracentrifugation through a 20% (w/v) sucrose cushion. The pellet was resuspended in PBS, snap-frozen and stored at −80 °C. Virions were incubated in presence of 700 nM LEN for 1.5 h at room temperature and then mixed with 10 nm colloidal gold beads (10 nm diameter). The sample solution was applied to a C-Flat 2/2 3C grid that had been glow-discharged prior to use. Grids were blotted and plunge-frozen in liquid ethane using an FEI Vitrobot Mark II, operated at 16 °C and 100% humidity. Tomographic tilt series were acquired on a TF2 Tecnai F20 transmission electron microscope under low-dose conditions at 200 kV, using a Falcon III direct electron detector. Data were collected at a magnification of 50,000×, with tilt increments of 3° over a range from −40° to +40°, and defocus values between −3 µm and −6 µm. Tomograms were reconstructed using the IMOD software package (version 4.9.0). Alignment of 2D projection images was performed using gold fiducial markers, and 3D volumes were generated by back projection of the tilt series.

## Author contributions

Conceptualization: GAV, TB and MG. Investigation: MG, CW and NR, Supervision: GAV. Writing - original draft: GAV, TB, CW and MG. Writing - review and editing: All authors. Methodology: MG, CW and NR, Resources: GAV and TB, Data curation: All authors, Validation: All authors, Formal analysis: All authors, Software: All authors, Project administration: GAV, Visualization: All authors, Funding acquisition: GAV.

## Data and materials availability

All data needed to evaluate the conclusions in the paper are present in the paper and/or the Supplementary Materials. Additional data related to this paper may be requested from the authors.

