## Supporting Information for "Mechanistic Insights into Lenacapavir-Induced Off-Pathway HIV-1 Capsid Assembly"

The PDF file contains:

Figs. S1 to S3
Movies S1 to S2

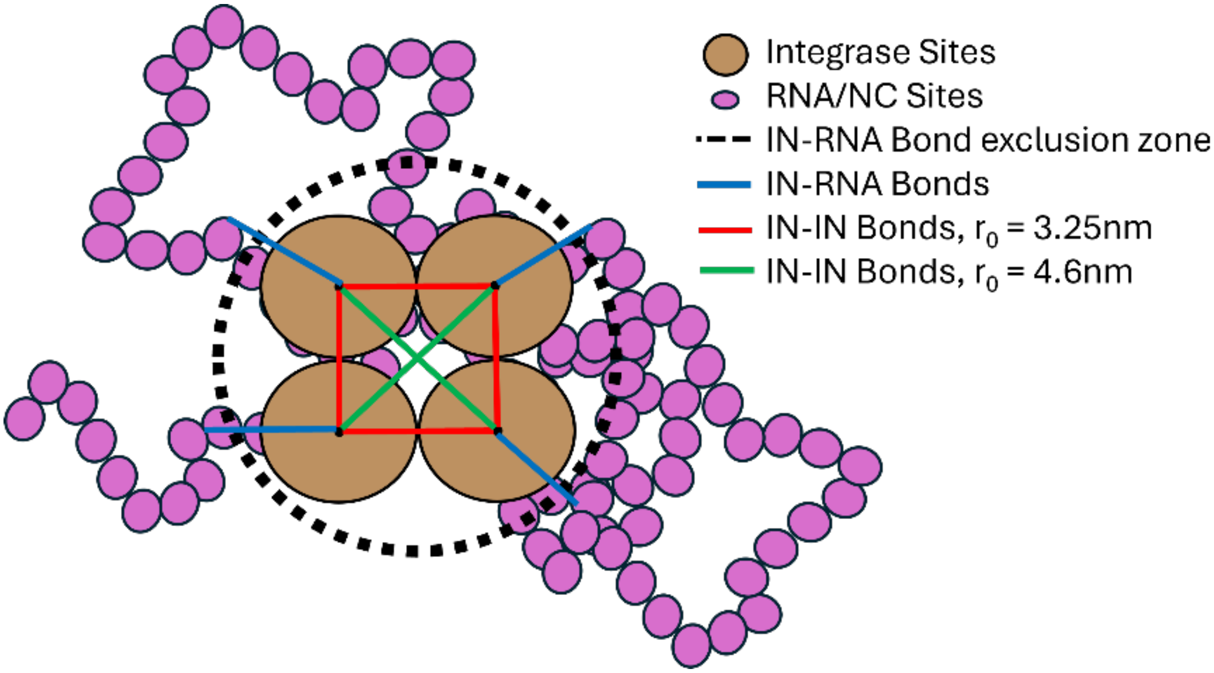


**Fig. S1. Model for the bonded topology of integrase crosslinking the RNA.**
As mentioned in the main text, the integrase tetramers consist of 4 beads bonded together, with each representing one integrase unit. The bond topology is shown in Figure X, where the red lines represent bonds where $r_0$ is 3.25nm and the green lines represent bond which are 4.6 nm long. Integrase units are also bonded to the RNA/NC beads. Using the first model with integrase tetramers and self-avoiding gag from Goodsell et al. as the initial structure, each individual integrase is bonded to the RNA/NC bead they are closest to, not including beads which are less than 4.6 nm from the center of the tetramer. This ensures that no RNA/NC bead is bound to more than integrase unit. The equilibrium distance for these bonds is set to the distance in the initial structure.

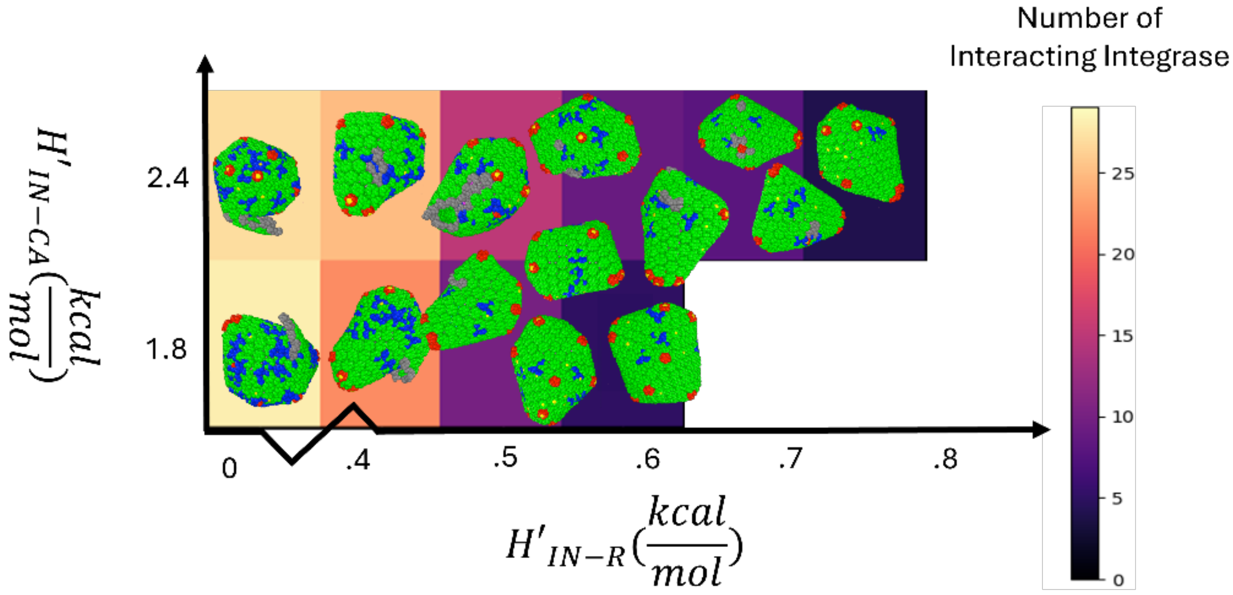


**Fig. S2. Assembly results in the absence of LEN.** different values of the integrase-RNP interaction strength, $H'_{\text{IN-R}}$ and integrase-CA interaction strength, $H'_{\text{IN-CA}}$. CA are shown.

The two main parameters that control the assembly of CA on the RNP are the interactions between CA and integrase (IN) and the interactions between IN and the rest of the RNP (R). CA-IN interactions control the amount of IN that is on the surface of the RNP and thus accessible to the CA, which will interact with the IN with a certain strength. In Figure Y, we tested various combinations of these parameters with only IP6 and no LEN present showing the capsids that encapsulate the RNP, if any. We also show the number of integrases that end up interacting with CA. We found that too many integrases interacting too strongly with the capsid led to defective capsids while not enough integrases interacting too weakly led to failed encapsulation. We choose the following values to balance these factors. $H'(IN-R)$ = 0.56 $kcal/mol$ and $H'(IN-CA)$ = 1.8 $kcal/mol$ correspond to the values of H given in the main text.

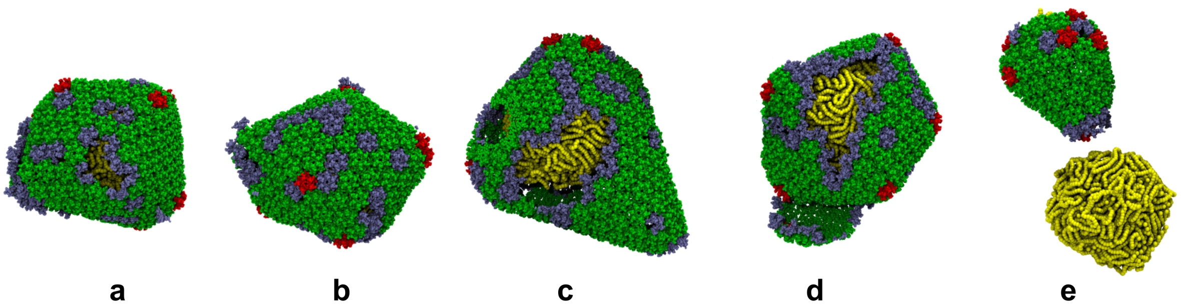

a b c d e

**Fig. S3. Additional simulations of capsid-RNP packaging in the presence of LEN**. (**a-e**) Simulation snapshots showing that capsid assembled malformed architectures in the presence of LEN. On few occasions LEN treated capsid failed to enclose RNP partially or completely.

**Movie S1: Capsid morphogenesis at low LEN levels.** All four independent trajectories demonstrate that capsid can adopt conical geometry, with pentamers forming during both early and late stages of assembly. CA assembly is like WT, with the broad end forming first and growing into a hook-like structure. However, CA lattice closure is delayed and substantial defects emerge at the high curvature regions.

**Movie S2: Capsid morphogenesis at high LEN levels.** In one trajectory, a partially conical capsid forms, while in others, the CA lattice grows rapidly from both ends and anneals through end-to-end joining, characteristic of tubular structures. Pentamers fail to form during the late stages, preventing proper lattice closure. In all cases, an additional CA layer assembles on the lattice, reflecting elevated assembly kinetics at high LEN concentrations.
